# The role and mechanism of NRF2 in endothelial cells and smooth muscle cells of pulmonary artery

**DOI:** 10.1101/2022.08.08.503173

**Authors:** Shasha Ning, Xinyue Guo, Yanan Zhu, Chenghui Li, Ruixue Li, Yinan Meng, Weiwei Luo, Dezhang Lu, Yupeng Yin

## Abstract

Pulmonary hypertension (PH) is a kind of fatal disease. There are no existing drugs that could reverse the diseas. NF-E2-related factor 2 (NRF2) is one of the most important moleculars in the range of cell protection. We examined the expression of NRF2 in PH models and explored the role of NRF2 in regulating abnormal phenotypes in pulmonary artery cells. We determined the expression level of NRF2 in lung tissues of PH model decreased significantly. We found that NRF2 was reduced in rat pulmonary artery endothelial cells (rPAEC) under hypoxia, while it was overexpressed in rat pulmonary artery smooth muscle cells (rPASMC) under hypoxia. Next, the results showed that knockdown NRF2 in rPAEC promoted endothelial-mesenchymal transformation and upregulated reactive oxygen species level. After the rPASMC was treated with siRNA or activator, we found that NRF2 could promote cell proliferation by regulating PDGFR/ERK1/2 and mTOR/P70S6K pathways, and accelerate cell migration by affecting MMP2/3/7. Therefore, the study suggests that the combination of NRF2 activators should be considered to eliminate the promoting effect of NRF2 activators on proliferation and migration of rPASMC.

**Summary statement:** Hypoxia regulates NRF2 in PAECs and PASMCs differently. We found that NRF2 could promote cell proliferation by regulating PDGFR/ERK1/2 and mTOR/P70S6K pathways, and accelerate cell migration by affecting MMP2/3/7.

## Introduction

Pulmonary Hypertension (PH), determined by an increase in mPAP (≥25mmHg at rest) as measured through a right ventricular catheter, could overload the right ventricle, leading to right ventricular failure and death ultimately (Mandras *et al*., 2020). The disease often appears progressive shrinkage of small pulmonary vessels and gradual occlusiveness of a pulmonary artery, (Tuder, 2017) which is mainly due to pathologic thickening of pulmonary vascular intima and tunica media vasorum (Thenappan *et al*., 2018). It is well known that endothelial dysfunction is the initial factor of pulmonary vascular remodeling (Derrett-Smith *et al*., 2013). Under normal physiological conditions, pulmonary vascular endothelial cells (PAEC), as an important barrier in blood vessels, play an important role in controlling permeability, regulating inflammation, balancing coagulation, and guiding angiogenesis (Dalal *et al*., 2020). However, in pulmonary hypertension, the pulmonary artery endothelial cells affected by inflammatory factors and angiogenic factors, and then it appears some abnormal phenotypes, such as endothelial dysfunction, excessive proliferation, resistance to apoptosis and Endothelial-mesenchymal transformation (EndMT) (Florentin *et al*., 2022). Those abnormal phenotypes could result in intimal thickening which promotes vascular remodeling. EndMT is a process in which endothelial cells are stimulated by TGF-β and then lose intercellular connections and change cell morphology to gain the capacities of proliferation and migration (Gong *et al*., 2020). At this time, cells express mesenchymal cell markers such as α-SMA and SM22 (Good *et al*., 2015). Existing studies have found that multiple signaling pathways are involved in EndMT process. The TGF-β/Smad pathway has been identified as a key pathway involved in EndMT (Wu *et al*., 2016). Smooth muscle cells are responsible for the contraction and relaxation of blood vessels and regulate blood flow to tissues and organs (Gong *et al*., 2020). In pathological conditions, smooth muscle cells show excessive proliferation, migration, and secretory activity while their contraction capacity was reduced (Wall and Bornfeldt, 2014). The proliferation and migration of pulmonary artery smooth muscle cells (PASMC) in pulmonary hypertension is the central events of pulmonary vascular remodeling, and the exploration of related aspects has important significance for the PH process and drug development. Current studies show that PDGF is an important signaling pathway regulating smooth muscle cell phenotype transformation, PDGFR inhibitors (imatinib) have a certain inhibitory effect on pulmonary hypertension (Mucke, 2013).

NF-E2-related factor 2 (NRF2) is a key redox-sensitive transcription factor in cells, which can enter the nucleus and regulate the transcription of downstream genes by binding to the ARE sequence of the gene promoter region, therefore it is capable of regulating the expression of redox-related enzymes and phase II detoxification enzymes to relieve the oxidative stress, inflammation and promotes cell survival (Kaspar *et al*., 2009). NRF2 can regulate cancer progression by affecting the proliferation, migration of cancer cells, and angiogenesis (Murakami and Motohashi, 2015)(Zhang *et al*., 2019; Huang *et al*., 2021) (Gao *et al*., 2021). Although activation of NRF2 has been found in some cancer cells to increase the antioxidant capacity and detoxification abilities of cells, continued activation of NRF2 in normal cells has a harmful effect (Kitamura and Motohashi, 2018). The dark side of NRF2 has been widely reported. For example, the overactivation of NRF2 plays an important role in the tumorigenic ability of cancer cells, such as their resistance to treatment and greater invasiveness (Paramasivan *et al*., 2019). In pulmonary hypertension, it has been found that activated NRF2 in endothelial cells significantly alleviated endothelial dysfunction (Chen *et al*., 2017). Other studies have found that NRF2 activation eased oxidative stress level of endothelial cells which is an important inducing factor of endothelial various abnormal phenotypes (Zhang *et al*., 2021). It has been found in smooth muscle cells that some miRNAs can regulate the proliferation and migration of smooth muscle cells by targeting Keap1/NRF2(Zhang *et al*., 2020) The NF-E2-associated factor 2 (NRF2)/ Kelch-like Ech-associated protein 1 (Keap1) system is a key component of the homeostasis oxidative stress response and plays an important role in neointimal hyperplasia(Ashino *et al*., 2016).

The important role of NRF2 in abnormal phenotypes of PAECs and PASMCs in PH has not been clearly explained. The purpose of this study was to explore the regulation and mechanism of NRF2 on various abnormal phenotypes of PAECs and PASMCs in the progression of pulmonary hypertension, increase the understanding of abnormal phenotypes of the two types of cells, and lay a theoretical foundation for the development of NRF2-related drugs.

## Results

### The expression of NRF2 in lung tissue of PH rat model

To explore whether NRF2 is involved in the disease progression of PH, we established MCT-PH and H-PH rat models. And right ventricular pressure was measured by catheterization. We found that right ventricular systolic pressure was significantly higher in MCT-PH and H-PH rats than in the control rats which means the disease model has been established successfully (Fig1 A-B). After that, the expression of NRF2 was detected in the lung tissues of control group and two types of PH model rats, and it was found that the protein and mRNA expression levels of NRF2 in the MCT-PH and H-PH rat lung tissues were significantly decreased (Fig1 C-F), and the protein expression levels of NRF2 downstream protein HO-1 in the H-PH rat lung tissue were significantly decreased (Fig1 E). In the meanwhile, the overall fluorescence intensity of NRF2 was significantly decreased which showed by immunofluorescence (Fig1 G-H). This indicated that NRF2 expression level was significantly inhibited in the lung tissues of MCT-PH and H-PH rat models.

**Fig. 1.**
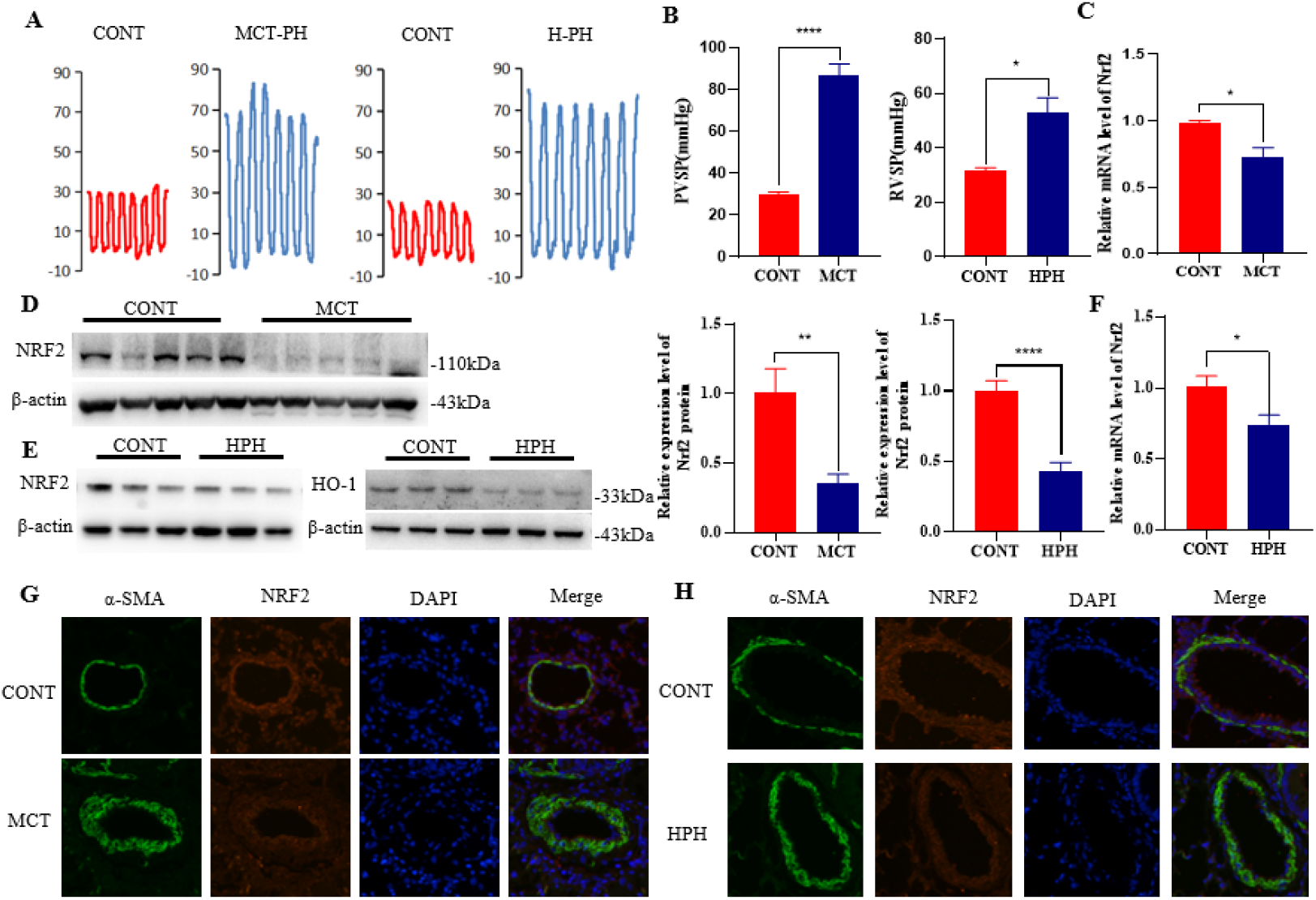
The expression level of NRF2 in MCT-PH and H-PH rat lung tissue. (A) Representative diagrams of right ventricular pressure of rats in the control, MCT-PH and H-PH groups. (B) Statistical analysis of right ventricular systolic pressure of the control group, MCT-PH and H-PH rats. <u>(C-F) The expression of NRF2 in lung tissues of control group and MCT-PH and H-PH model group. (G-H) Representative immunofluorescence images of NRF2 in lung tissues of the control group and MCT-PH model group.</u>

### The effect of NRF2 on EndMT of rPAEC

During pulmonary hypertension, pulmonary vascular cells are under hypoxia, which activates the hypoxia-inducible factor protein and triggers multiple downstream reactions. To explore the effect of hypoxia on NRF2 expression in PAECs, we detected the expression level of NRF2 and its downstream protein HO-1 by western blot in ECs treated with hypoxia. The results showed that NRF2 and HO-1 protein level is down-regulated both in hPAEC and rPAEC treated with hypoxia (Fig2 A-B). And the mRNA expression level of NRF2 in rPAEC under hypoxia is also decreased (Fig2 C). Then, we knocked down NRF2 to explore its effect on EndMT. Firstly, the efficiency of three siRNA was detected under normoxic conditions, and si-558 was selected for the following experiments according to western blot results (Fig2 D). Next, the rPAEC was treated with TGF-β1(EndMT inducer) for 4d after when NRF2 was knocked down under normoxic conditions. The results showed that α-SMA/SM22 (the markers of mesenchymal cells) increased significantly after 4 days of TGF-β1 treatment. Importantly, mesenchymal markers of NRF2 knockdown combined with TGF-β1 treatment group were significantly and further increased compared with TGF-β1 alone (Fig2 F). This suggests that NRF2 knockdown promotes EndMT phenotype and may imply that NRF2 plays a palliative role in EndMT. To explore its mechanism, NRF2 was knocked down under normoxic and hypoxic conditions to study its influence on ROS which is one of the most important inducers of EndMT. It was found that the total ROS level in rPAEC increased significantly after the knockdown of NRF2 (Fig2 E). The result confirmed the regulation of ROS by NRF2 in rPAEC. And we also detected the changes of Snail1 protein level after NRF2 knockdown which is an important transcription factor of EndMT. We found that Snail1 protein levels were significantly increased in the TGF-β1-treated group compared with the control group. And compared with TGF-β1 alone, Snail1 protein levels were significantly increased in NRF2 knockout combined with TGF-β1 treatment group (Fig2 F). Therefore, NRF2 might regulate EndMT by influencing the protein level of Snail1 and ROS levels in PAECs.

**Fig. 2.**
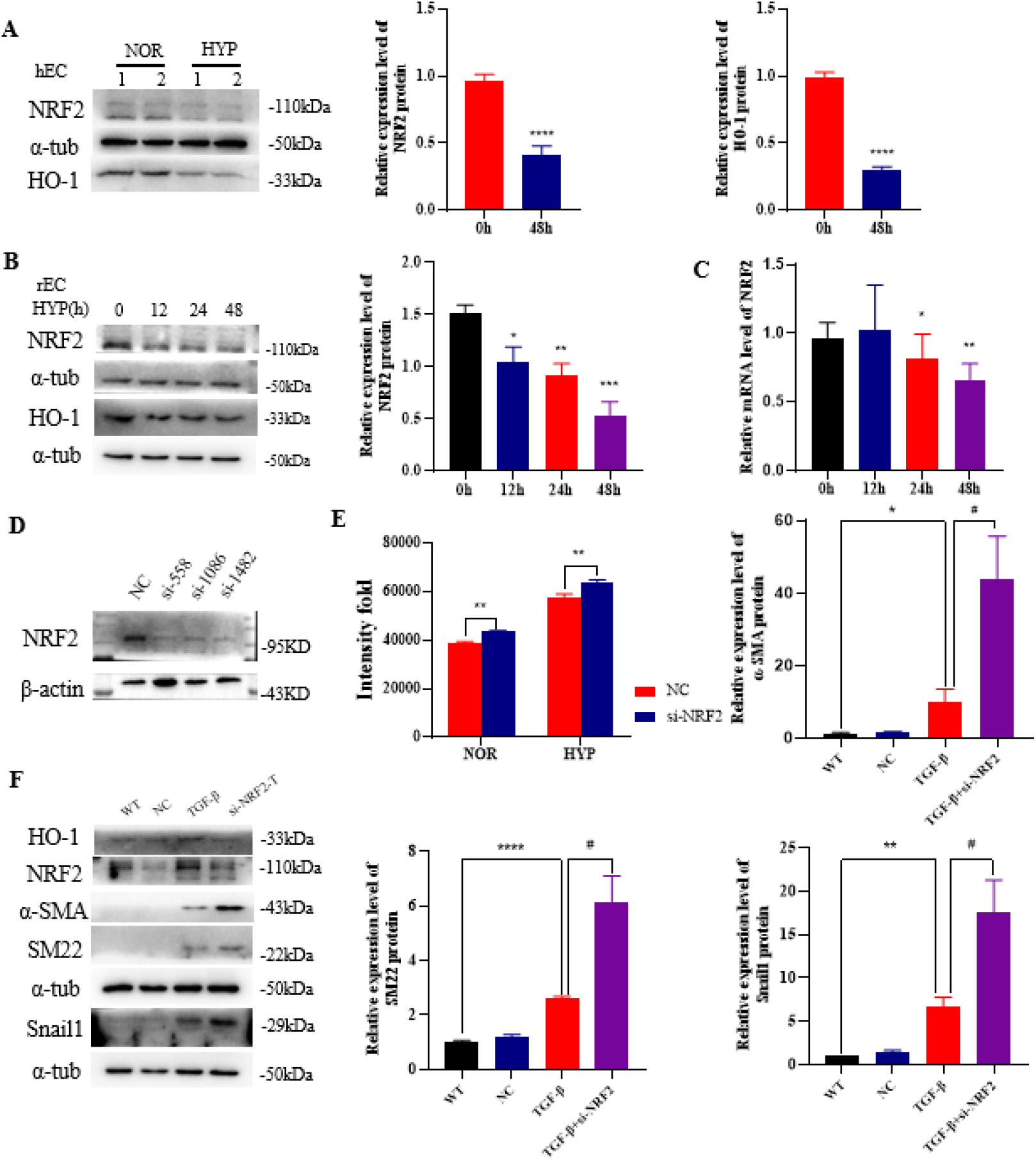
The effect of NRF2 on EndMT in rPAEC. (A-B) hPAEC and rPAEC were treated with hypoxia, western blot was used to determine NRF2/HO-1 protein expression changes and quantitative analysis was performed. (C) The mRNA expression level of NRF2 in rPAEC under hypoxia was detected by RT-qPCR. (D) After treatment with si-NRF2 in rPAEC, the interference efficiency was measured by western blot assay. (E) After treated with si-NRF2 in rPAEC, the total ROS level was detected by ROS kit. (F) rPAEC were treated with TGF-β1 after si-NRF2, western blot was used to detect EndMT marker(α-SMA,SM22) and Snail1, and quantitative analysis was performed.

### The regulation of NRF2 on the proliferation of rPASMC

To explore the role of NRF2 in PASMC, we detected the expression level of NRF2 after hypoxia treatment. We found that NRF2/HO-1 protein level is up-regulated in hPASMC and rPASMC under hypoxia which is different from ECs (Fig3 A-B). The result may suggest that NRF2/HO-1 is required for the maintenance of cellular physiological functions in highly proliferative smooth muscle cells. It has also been described that smooth muscle cells with high proliferation may be in a state of “NRF2 addiction”. Next, we knocked down NRF2 to explore its effect on rPASMC proliferation. Firstly, NRF2 was knocked down under normoxic and hypoxic conditions, and the changes of PCNA protein expression level were detected by western blot. Our results showed that knocking down NRF2 in normoxic and hypoxic states could significantly inhibit the protein level of PCNA (Fig3 C). In the meanwhile, the results showed that NRF2 knockdown significantly upregulated α-SMA protein (Fig3 C). This suggests that silencing NRF2 can alleviate the loss of smooth muscle contraction phenotype during the disease. Subsequently, EdU and CCK experiments were carried out to verify the above experimental results, and the results showed that knocking down NRF2 under normoxic and hypoxic states could significantly inhibit the proliferation level of smooth muscle cells (Fig3 D-E). In conclusion, NRF2 knockdown can significantly inhibit rPASMC proliferation.

**Fig. 3.**
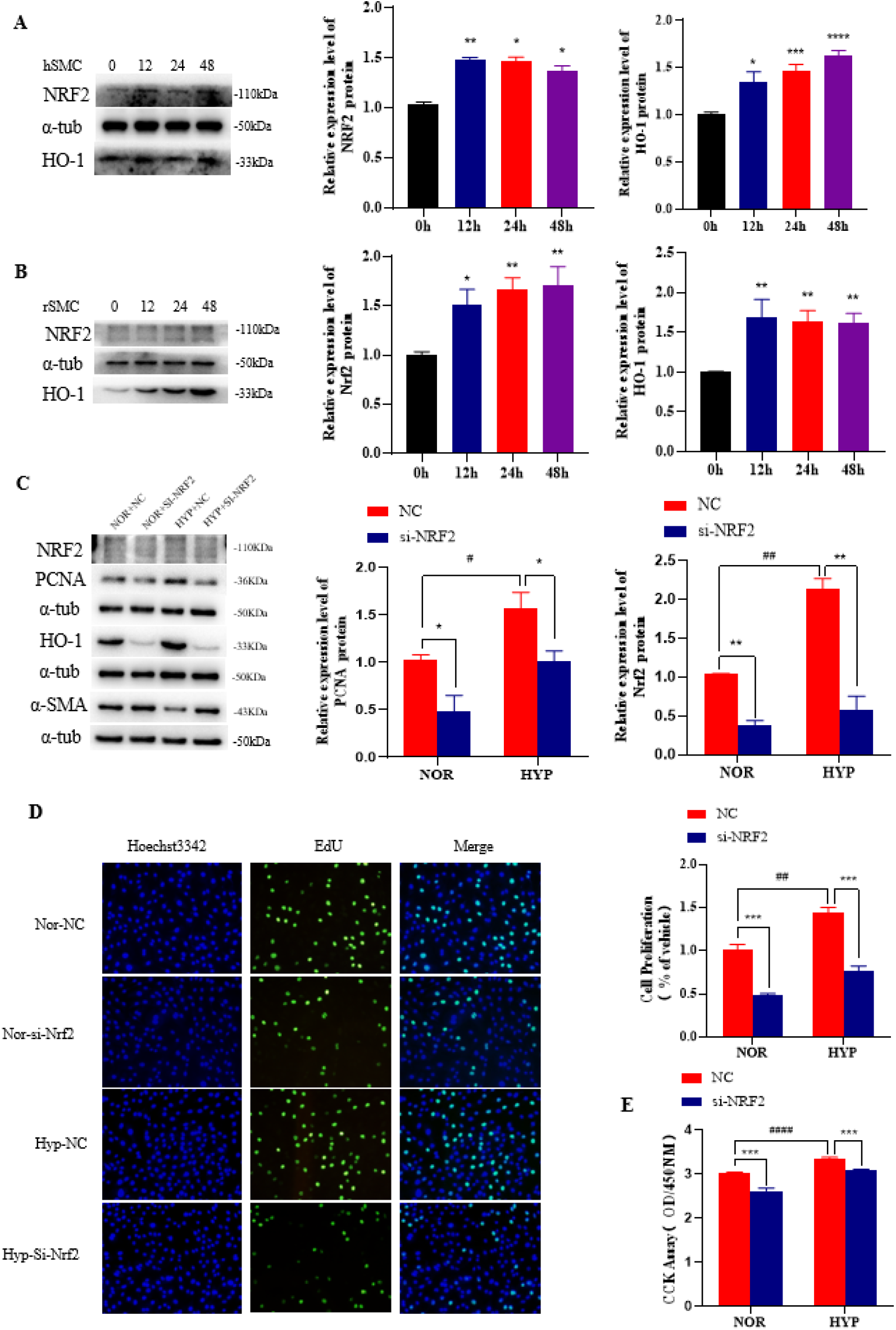
The effect of NRF2 on proliferation of rPASMC. (A-B) The expression of NRF2/HO-1 was detected by western blot after treated with hypoxia in hPASMC and rPASMC. (C) Knockdown of NRF2 was performed under normoxic and hypoxic conditions, and the expression of PCNA/α-SMA protein was detected by western blot and quantitative analysis was completed; (D) Knockdown of NRF2 was performed under normoxic and hypoxic conditions, EdU was completed to detect cell proliferation; (E) Knockdown of NRF2 was performed under normoxic and hypoxic conditions, CCK8 were completed to detect cell proliferation.

### The regulation of NRF2 on the mTOR/P70S6K and PDGFR/ERK1/2 pathways

In order to further determine the regulation of NRF2 on rPASMC proliferation, we treated rPASMC with NRF2 activator SFP under normoxic and hypoxic conditions to explore the effect of SFP on rPASMC proliferation through CCK experiment. Firstly, cells were treated with different SFP concentrations to find the appropriate effect concentration. The result shows that 5uM SFP had a higher activation level of NRF2/HO-1 and a higher stimulation effect on PDGFR-α protein which is an important protein that we found NRF2 could regulate, so 5uM was selected as the final action concentration (Fig4 A). During the experiment, we found that high concentration of SFP (≥20uM) has a greater capability of killing cells. Next, NRF2 activation were treated under normoxic and hypoxic conditions, and the CCK test result shows that the rPASMC proliferation level increased significantly after activation under normoxia and hypoxia (Fig4 B).

**Fig. 4.**
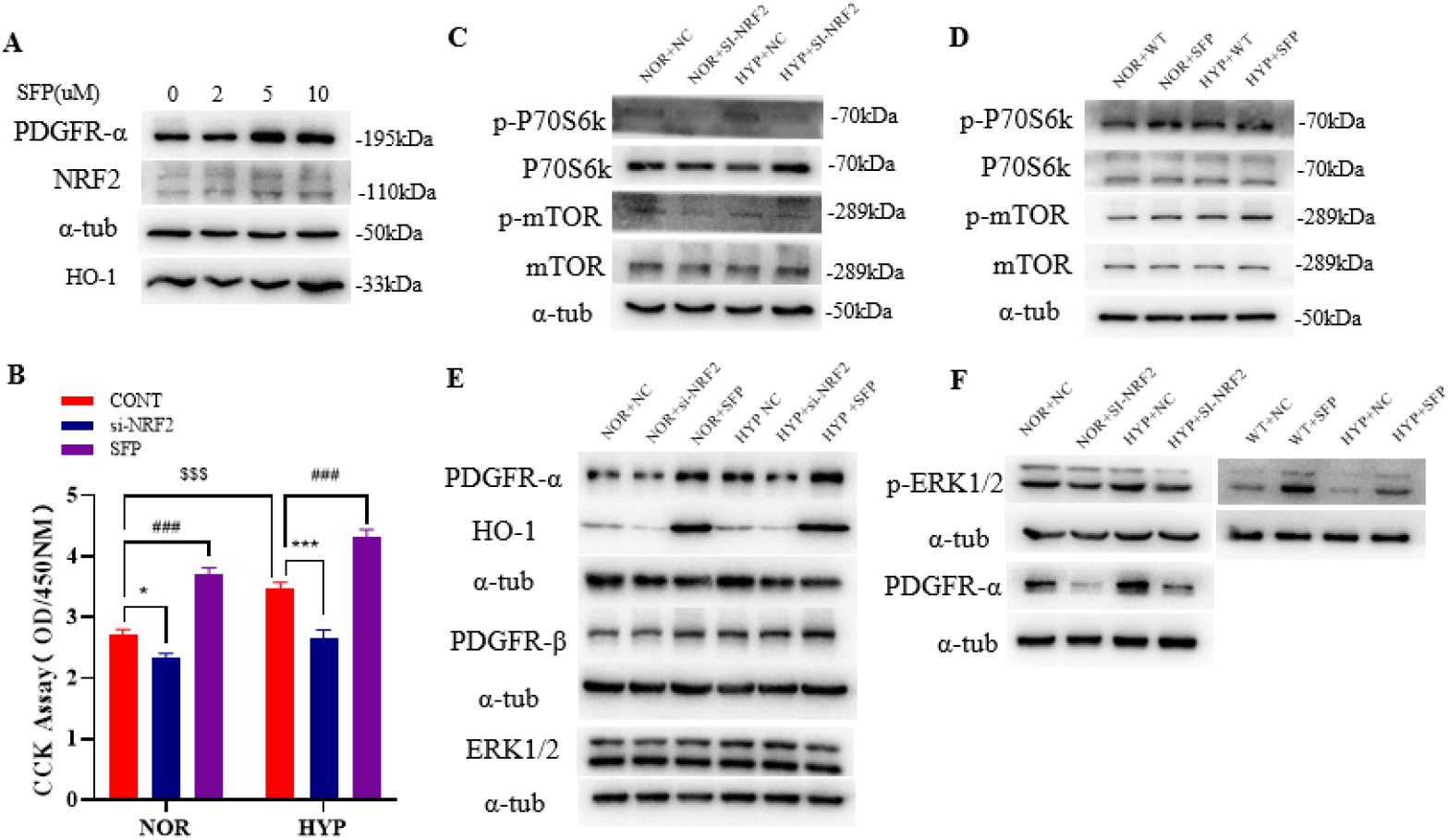
The regulation of NRF2 on mTOR/P70S6k and PDGFR/ERK1/2. (A) rPASMC was treated with SFP gradient concentration, and the activation of NRF2 was detected by WB. (B) rPASMC was treated with 5uM SFP in normoxic and hypoxic conditions and cell proliferation was measured by CCK8. (C-D) NRF2 was knocked down and activated under normoxic and hypoxic conditions, and the relative protein levels of p-mTOR /mTOR and p-P70S6K/P70S6K were detected by Western blot. (E-F) NRF2 was knocked down and activated under normoxic and hypoxic conditions, and the protein levels of PDGFR-α and ERK1/2/p-ERK1/2 were detected by western blot.

To explore the specific mechanism of NRF2 regulating cell proliferation, we studied one of the most important pathways mTOR/P70S6K. mTOR /P70S6K has been shown to play an important role in cell proliferation and migration. We knocked down NRF2 in normoxic and hypoxic states, and we found that compared with si-NC group, the protein level of phosphorylated mTOR/P70S6K in the NRF2 knockdown group was significantly decreased (Fig4 C). To further verify the regulation of NRF2 on this pathway, we detected the protein levels of phosphorylated mTOR and P70S6K after SFP treatment, and we found that the protein levels of phosphorylated mTOR and P70S6K increased significantly after SFP treatment (Fig4 D).

PDGF is an important molecule that regulates the phenotypic changes of pulmonary arterial smooth muscle, the proliferation and migration of PASMC(Humbert *et al*., 2004). To explore the regulation of NRF2 on PDGF in pulmonary arterial smooth muscle cells, protein levels of PDGFR and ERK1/2 were detected after NRF2 knockdown and activation. The results showed that PDGFR-α protein level decreased significantly after NRF2 knockdown (Fig4 E-F). Therefore, we examined the presence of ARE sequences in the PDGFR promoter region, and we found that both PDGFR-α and PDGFR-β had more than one ARE binding sequence which partially explains the reason why NRF2 regulates the PDFGR. At the same time, the protein level of P-ERK1/2 showed a significant decrease after NRF2 knockdown. And it was further activated after being treated with SFP (Fig4 E-F). These results confirmed the regulation of NRF2 on mTOR/P70S6K and PDGFR/ERK1/2 in PASMC.

### The regulation of NRF2 on migration of smooth muscle cell

To investigate the regulation of NRF2 on the migration of rPASMC, we treated rPASMC with si-NRF2 and SFP in normoxic and hypoxic states, respectively. We found that knocking down NRF2 in both normoxic and hypoxic states could significantly inhibit the migration of rPASMC (Fig5 A). After SFP treatment, the migration ability of rPASMC was significantly enhanced (Fig5 E). To further explore the basis of its action, matrix metalloproteinases (MMP2/3/7) associated with cell migration were detected. We found that mRNA levels of MMP2/3/7 increased significantly after hypoxia compared to control. And compared with si-NC group, NRF2 knockdown significantly inhibited the mRNA levels of MMP2/3/7 (Fig5 B-D), and the mRNA levels of the above proteins increased significantly after NRF2 activation (Fig5 F-H).

**Fig. 5.**
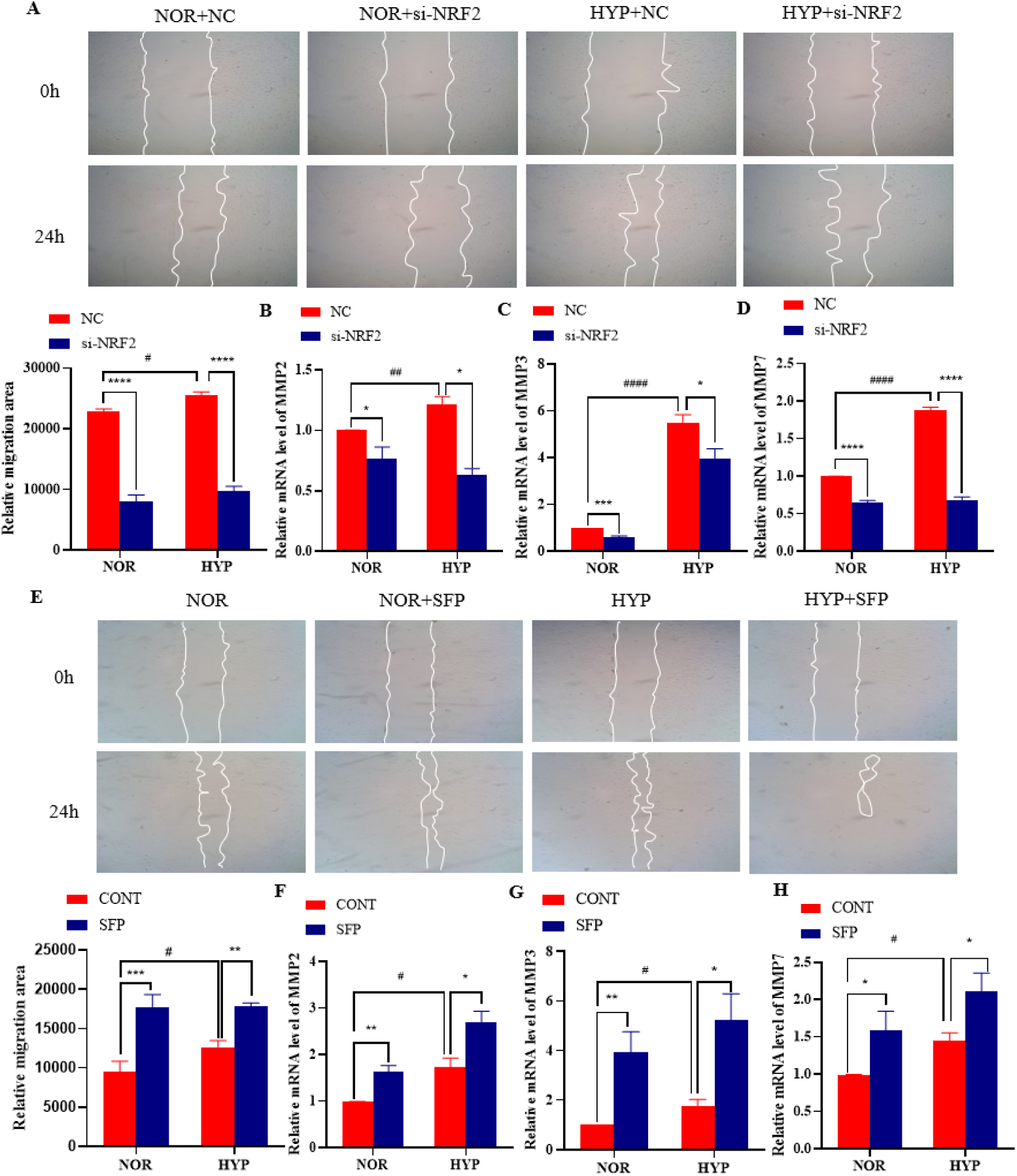
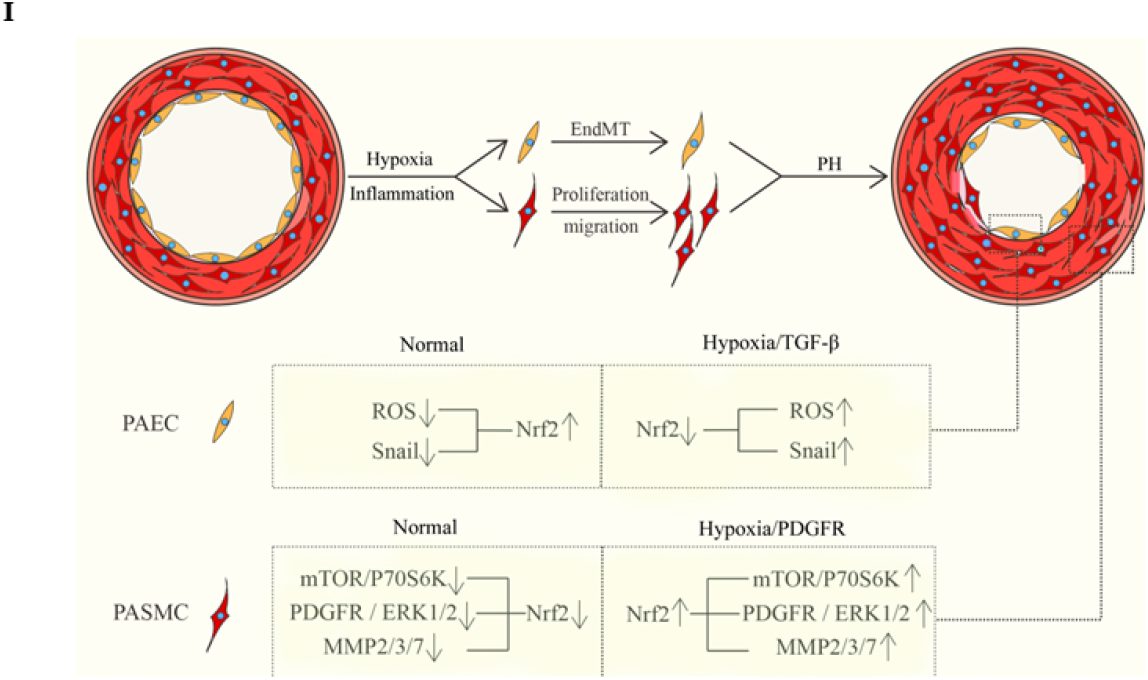
The effect of NRF2 on migration of rPASMC. (A) NRF2 in rPASMC was knocked down under normoxic and hypoxic conditions, and the cell migration level was detected by scratch test. (B-D) NRF2 was knocked down in rPASMC under normoxic and hypoxic conditions, and the expression of MMP2/3/7 was detected by RT-qPCR. (E) NRF2 in rPASMC was treated with SFP under normoxic and hypoxic conditions, and the cell migration level was detected by scratch test. (F-H) NRF2 was treated with SFP in rPASMC under normoxic and hypoxic conditions, and the expression of MMP2/3/7 was detected by RT-qPCR. (I) NRF2 knockdown inhibited the proliferation and migration of rPASMC by down-regulating PDGFR/ERK1/2, mTOR/P70S6K and MMP2/3/7. These results indicated that NRF2 plays different roles in PAEC and PASMC.

## Discuss

During pulmonary hypertension, PAECs and PASMCs exhibit various abnormal phenotypes that are responsible for promoting disease progression. Currently, the knowledge and understanding of these abnormal phenotypes is not sufficient. This study attempted to explore the regulation of NRF2, an important transcription factor in cells, on these abnormal phenotypes, and attempted to provide a research basis for the development of related drugs. It has been suggested that activation of NRF2 in inflammatory lung diseases may initiate protective mechanisms to mitigate disease progression. Existing studies have reported the therapeutic effect of NRF2 activator in pulmonary hypertension. In animal studies, the researcher found that NRF2 activator can partially alleviate the course of pulmonary hypertension.(Zhang *et al*., 2022) Meanwhile, the researcher suggested that the therapeutic effect of NRF2 might be achieved by alleviating lung inflammation in SuHx-PH model. Unfortunately, the researchers did not perform a cell-level analysis. Other researchers confirmed some conclusions in animal experiments, and also found the effect of NRF2 activator on right heart injury and vascular remodeling(Kang *et al*., 2020). However, neither of them was explored at the cellular level, nor did they involve the regulatory effect of NRF2 on pulmonary vascular cells, and those researches mainly focus on the protective effect of NRF2 on right heart function. However, our study found that NRF2 plays different roles in PAECs and PASMCs, and has different regulatory roles in abnormal phenotypes of the two types of cells. Our results of NRF2 in ECs are consistent with those of another investigator (Chen *et al*., 2017b). These researchers not only found that NRF2 activator can alleviate MCT-PH lung inflammation, but also confirmed that NRF2 activation can improve the functions of PAECs. On this basis, our study supplemented the mechanism of the regulation of NRF2 on EndMT, that is, NRF2 affects EndMT by regulating the protein level of Snail1 and ROS level. Although activation of NRF2 appears to have a positive effect on the treatment of PH, our study in PASMC does not support this view. Our study found that NRF2 knockdown inhibited the proliferation and migration of rPASMC by down-regulating PDGFR/ERK1/2, mTOR/P70S6K and MMP2/3/7. These results indicated that NRF2 plays different roles in PAEC and PASMC. Correspondingly, hypoxia regulates NRF2 in PAECs and PASMCs differently. We found that hypoxia down-regulated NRF2 expression level in PAECs but up-regulated NRF2 expression level in PASMCs. This may be because over-proliferating PASMC are more similar to some cancer cells than PAEC and require antioxidant and other activities of NRF2 to maintain cellular homeostasis, or because of a large number of downstream genes of NRF2 lead to complex and multifaceted physiological activities. Combined with previous studies and our current results, we speculated that the effects of NRF2 on PH and pulmonary vascular remodeling may be achieved through complex regulation at multiple levels. However, due to the different roles of NRF2 in PAEC and PASMC, NRF2 activator cannot be used alone to treat the disease, and combination should be considered.

## Methods

### Animal model

Adult male Sprague Dawley rats purchased from Beijing Vital River Laboratory Animal Technology (Beijing, China), 180 to 200g body weight. All the rats were housed with food and water ad libitum, and lived in a temperature-controlled room at 20-25°C under a regular 12 h/12 h light/dark schedule. All of the animal experimental procedures were approved by the Institute of Animal Care and Use Committee at Chinese Academy of Medical Sciences.

For establishment of MCT-induced pulmonary hypertension model, the rats were randomly divided into control and MCT group. MCT group rats received a single intraperitoneal injection of 50 mg/kg monocrotaline (MCT; MedChemExpress; US). Dosage was calculated after weighing rats. MCT was dissolved in 1 M HCl at a concentration of 40 mg/ml, adjusted pH to 7.4 with 0.5 M NaOH solution and diluted with distilled water. And then after the 28-days normal feeding, rats were sacrificed for experimental measurements.

For hypoxia-induced pulmonary hypertension model, the rats were randomly divided into control and hypoxic group. The hypoxic rats were placed in an automatic hypoxic chamber. The hypoxic chamber was continuously filled with nitrogen to maintain the overall oxygen concentration at 10%. CO_2_ absorbent and desiccants were put into the hypoxic chamber to absorb exhaled CO_2_ and moisture for 4 weeks, and change it once a week. The control group of rats were bred in the same room under normoxia. After 28-days of feeding, rats were sacrificed for experimental measurements.

### Right ventricular pressure measurement

Rats were anesthetized with isoflurane by a ventilator. When rats lost reaction after pinching their toes, the neck of rats was sprayed with alcohol. Then the skin on the left side of the neck was cut of rats, use forceps blunt to separate the muscle and connective tissues. After exposure to the carotid artery, use small scissors to make an incision in the vein (not cut the whole blood vessel), and then insert a catheter at the incision and slowly push it to the right ventricle. At this time, attention should be paid to the pressure waveform on the computer connected to the instrument. When the waveform is stable enough, the actual right ventricular systolic and diastolic blood pressure should be recorded.

### Cell culture

Human pulmonary arterial endothelial cells and smooth muscle cells were obtained from Prof. Haiyang Tang of our laboratory. Rat arterial endothelial cells and smooth muscle cells were obtained from our laboratory. PAECs were maintained in Endothelial Basal Medium (EBM-2; Univ; Shanghai) supplemented with 10% fetal bovine serum. SMCs were maintained in Dulbecco’s Modification of Eagle’s Medium (DMEM; Procell; Wuhan) supplemented with 10% fetal bovine serum (FBS; Procell; Wuhan). Cells were cultured at 37 °C in a 5% CO_2_ incubator.

### Western blot

Lung tissue preserved in liquid nitrogen was removed, after the rough grinding in a precooled mortar, the tissue powder is placed in a grinding tube with protein lysis fluid. The grinding tube is then put into a high-throughput tissue grinder for secondary grinding. 80Hz, 1min/ time, repeat 5-6 times. After 15min ice lysis, centrifugation was performed at 12000g at 4°C for 10min. The supernatant was absorbed and the protein concentration of each sample was measured by bicinchoninic acid (BCA Kit; Beyotime Biotechnology; Shanghai). After calculation, protein lysis solution and 5X Loading Buffer were added respectively for protein dilution. Mix well and centrifuge it. Heated water bath to 100□ and boiled for 10 min to denaturant proteins.

Electrophoresis gel was prepared according to the standard. The protein bands were separated after loading and electrophoresis, and the protein was transferred to PVDF membrane. The membrane was sealed with 5% defatted milk powder for 1h, and the primary antibody was incubated at 4°C overnight. After receiving the primary antibody, TBST was washed three times, 10min/ time. The second antibody was incubated according to the source of the primary antibody, and TBST was washed three times, 10min per time. Membranes were exposed to a mixture of enhanced chemiluminescence solution and detected by Tanon automatic chemiluminescence imaging (Tanon, Shanghai). Quantitation of protein band was performed by Image J software.

### Real-time fluorescence quantitative PCR

The grinding part of RNA extraction is the same as that of protein extraction. Trizol kit was used to extract RNA from tissues after tissue grinding. Primers were designed and synthesized, and then PCR was performed to check whether the synthesized product was a band. The RT-qPCR reaction liquid system was prepared according to the primers 2ul, cDNA 2ul, RNase-free water 6ul and SYBR Green 10ul. With β-actin or 18S as reference, the two-step setting procedure is 95°C 30s, 95°C 5s, 60°C 30s, 40 cycles, which was running by Bio-rad CFX Connect Fluorescence quantitative PCR instrument (Bio-rad, US). The relative circulation number of the target gene was detected. The primer sequences of rat NRF2 were F: TTGCCGCTCAGaACTGTagg, R: GGaacaagGAACACgTTgcc.

### Immunofluorescence staining

During sampling, the same parts of lung tissue were fixed with 4% formaldehyde perfusion and placed on a shaker for 2d continuous fixation. After ethanol dehydration, infiltration, embedding and sectioning, blank tissue sections were obtained. The sections were observed under a light microscope before the immunostaining procedure. Paraffin sections were routinely dewaxed and dehydrated before antigen repair. It was brought in the microwave oven until it boils slightly, then stop heating immediately and repeat 3-4 times. PBST was infiltrated 3 times, 3min/ time. At room temperature, the permeability was carried out for 20min, and then wash it with PBST 3 times, 3min/ time. Configure 5% BSA and seal the sections with it for 15 minutes. After drying the sealing solution, the primary antibody was incubated at 4°C overnight or room temperature for 1h. PBST was infiltrated 3 times, 3min per time. The second antibody was incubated at room temperature for 1h, and then wash it with PBST 3 times, 3min/ time. It used reagent containing DAPI for final sealing tablets. Be careful to avoid slide movement after sealing.It was placed in a black box with water to avoid light and take to an inverted fluorescence microscope(Zeiss, Germany) for observation.

### CCK8 cell proliferation assay

After subculture, the cells were placed in a 96-well plate for culture. After the cell growth density reached 60-70%, the cells were performed NRF2 knock-down. After cultured for 24h, 10ul CCK8(ZETA LIFE; US) solution was added to each well without changing the liquid, and bubbles should not be generated. After incubation for 4h in the incubator, the absorbance value at 450nm was detected by a microplate reader.

### EdU cell proliferation assay

The cells were subcultured and placed in 96-well plates for culture. After the cell growth density reached 60-70%, the cells were performed NRF2 knock-down. After cultured for 24h, configure EdU(Beyotime; Shanghai) working fluid 2X according to the number of wells and preheat it at 37°C. EdU working solution was added in equal volume and incubated for 2h. The culture medium was removed and fixed at room temperature for 15min. The cells were washed 3 times after discarding the liquid, and the permeability liquid was added for permeability. The cells were washed 3 times after discarding the liquid, 3-5min/time. The reaction solution was formulated according to the instructions and evenly added to the cell surface. The cells were incubated with foil at room temperature, away from light. Wash 3 times after discarding liquid. Hochest33342 was diluted with PBS and incubated for 10 min at room temperature without light, then washed with the liquid 3 times. It can be observed and photographed under a positive fluorescence microscope.

### SFP processing cell

SFP (Abcam; Cambridge) liquid is soluble in DMSO with a solution of 100mM. In smooth muscle cells, NRF2 activation level was detected after treatment with a gradient concentrations of 1uM, 2uM, 5uM and 10uM for 48h. After selecting the active concentration of SFP, the subsequent experiment was carried out accordingly. CCK8 experiment is the same as 4.1.2.3, in which the interference is changed to SFP treatment.

### Scratch test

After the cells were transferred to a six-well plate, they were cultured in a CO_2_ incubator until the cell density reached 80%-90%, and the cells were treated with scratches. Firstly we prepared a piece of white paper, draw 5-6 parallel lines on the paper with the help of a ruler, and then draw a line perpendicular to the existing line. We put the white paper under the six-hole plate. At the same time, 5-6 straight lines are in the horizontal position. We draw a straight line perpendicular to the petri dish with the yellow spear head according to the last drawn one. After each hole is marked, discard the medium. After the cells were washed away with PBS, the same amount of new medium was added, and at the same time, NRF2 knockdown and SFP treatment were performed. After culturing for 24h, the six-well plate was taken under the microscope to observe and record the scratches.

## Data analysis and statistics

Data are shown as mean ± standard error of the mean (SEM). Prism 8.0.2 software was used to perform statistical analysis. All data were submitted to Unpaired t-test or one-way analysis of variance (ANOVA) followed by Dunnett’s multiple comparison test. Differences were considered statistically significant at p-values less than 0.05. Significance code presented in the figures: ns = no significance, \**p* < 0.5, \*\**p* < 0.01, *** *p* < 0.001, **** *p* <0.0001.

## Acknowledgements

We would like to thank Prof. Haiyang Tang for the human cells.

